# *Learn!Bio* - A time-limited cross-sectional study on biosciences students’ pathway to resilience during and post the Covid-19 pandemic at an UK university from 2020-2023 and insights into future teaching approaches

**DOI:** 10.1101/2024.03.06.583815

**Authors:** Katy Andrews, Rosalie Stoneley, Katja Eckl

## Abstract

Higher education in biosciences is significantly informed by hands-on field trips and practical laboratory skills-training. With the first Covid-19 national lock-down in England in March 2020, on-campus education at higher education institutions was swiftly moved to alternative provisions, including online only options, a mix of synchronous or asynchronous blended, or hybrid adaptions. Students enrolled on an undergraduate bioscience programme have been faced with unprecedented changes and interruptions to their education. This study aimed to evaluate bioscience students’ ability to adjust to a fast-evolving learning environment and to capture students’ journey building up resilience and graduate attributes.

Bioscience undergraduate students in years 1-3 at the biology department at a Northwest English university participated in this anonymous, cross-sectional, mixed-method study with open and closed questions evaluating their perception and feedback to remote and blended learning provisions during the Covid-19 pandemic and post pandemic learning capturing academic years 2019/20 to 2022/23.

The Covid-19 pandemic and the consequent restriction of personal social interaction resulted in an significant decrease in the mental wellbeing of undergraduate bioscience students in this study, cumulating in poor or very poor self-rating of wellbeing in spring 2021; while at the same time students showed evidence of advanced adaption to the new learning and social environment by acquisition of additional technical, social and professional graduate-level skills, indicative of an, albeit unconscious, transition to resilience. Post pandemic, bioscience students worry about the increased living costs and are strongly in favour of a mixture of face-to-face and blended learning approaches.

Our results show that bioscience students can experience poor mental health while developing resilience, indicating tailored support can aid students’ resilience performance. Students have adjusted with ease to digital teaching provisions and expect higher education institutions continue to offer both, face-to-face, and blended teaching, reducing the burden on students’ significantly risen living costs.

## Introduction

In March 2020 the Word-Health Organisation (WHO) declared Covid-19 a pandemic [1]. In the UK, the government legislated the first national lockdown, which commenced March 26^th^, 2020, resulting in a public standstill and a closure of all face-to-face and on-campus teaching at English universities [2, 3] with most practical courses, laboratory experiments and field excursions suspended or converted to online or blended provisions for the majority of academic years 2019/2020 and 2020/2021 [4], respectively.

Benchmark statements for bioscience students’ education (2023) outlined by the Quality Assurance Agency for Higher Education (QAA) highlight the need for practical skills training in the laboratory, field, and computer/IT settings. During their studies, bioscience students can also expect to experience “*authentic research*” throughout their course while acquiring competencies in the lab, field, and computational practical and transferable skills essential for a post-graduate role [5]. A paper presented by Coward and Gray (2014) for the Royal Society for Biology has calculated an average bioscience student in the UK can expect 500 hours of practical training in their three years as an undergraduate student, while speciality training in the laboratory, field, or computational education might depend on the chosen specific subject [6].

The Covid-19 pandemic restricted students’ access to laboratories and research facilities, and most field trips and excursions had been cancelled. HEI providers sought alternative strategies to prepare students for a successful post-graduation career, including access to interactive online lab sessions, the introduction of lab or field virtual simulations, and lab/field activities to be conducted at home with any required material sent by post to all participating students [7–10].

With student accommodations closed and campus and social interactions significant reduced [11, 12], the Covid-19 pandemic has changed higher education teaching and learning world-wide, and has impacted on students’ wellbeing and their pathway to personal and career-related resilience and skills development in a yet not fully understood extend [13, 14].

This manuscript presents students’ feedback and voices from 317 participants surveyed from 2020-2023 in a longitudinal, cross-sectional mixed-methods study conducted at a North English university on bioscience students’ perception on their higher education during and post the Covid-19 pandemic and how the closure of laboratories and limitations to access campus facilities impacted their mental wellbeing and pathway to resilience development. Follow up surveys conducted in 2022 and 2023 post-pandemic compares students’ perception on the consequences of the pandemic on bioscience learning and career progression.

We hypothesised (1) most bioscience students exhibit the ability to adjust to a fast-evolving learning environment and build up resilience. We also hypothesised (2) that students preferred to return to a pre-pandemic mode of teaching. Post-pandemic, students are significantly supportive of a blended teaching concept but are not in favour of purely online learning.

## Methods

### Study design and study setting

This anonymous, cross-sectional, mixed-method cohort study followed a convergent parallel design with a focus on quantitative data collection and analysis. The study captured students’ feedback at four time points between November 2020 and May 2023 evaluating perception and acceptance of changes to **(1)** Learning during the pandemic & adjusting to a new challenging situation **(2)** wellbeing & resilience, and **(3)** consequences of the pandemic & students’ expectations from providers as reported by participants [15]. All participants have been in education during the Covid-19 pandemic, either at school or university level.

All students enrolled in an undergraduate bioscience degree (years 1-3) at Edge Hill University were eligible to participate in this study. Students were invited by email which entailed a direct link to the respective JISC online survey studies. Each study was accessible on the JISC online survey platform for four weeks, with one email reminder sent two weeks after the first invitation. Participation to each study was entirely voluntary and not linked to the participation in a previous or follow-up study. We completed four studies: Study-survey S1 (November 2020, mid-pandemic I, data collected from 23/08/2020 to 08/12/2020), study-survey S2 (April 2021, mid-pandemic II, data collected from 28/04/2021 to 04/05/2021), study-survey S3 (March 2022, follow-up I, data collection from 08/03/22 at 9am to 06/04/22), and study-survey S4 (May 2023, follow-up II, data collection from 03/04/2023 to 02/05/2023). All participants in this study were 18 years of age or older and fully able to consent.

### Student cohort and participants

Cohort sizes and number of participants for each study are shown in **table 1**. No personal identifiers were collected.

**Table 1:** Learn!Bio longitudinal, cross-sectional study 2020-2023: Cohort sizes and participants – overview.

| **Study** | **Date** | **Cohort size\*** | **Participants** | **Survey length** | **Completion**<br>**time** |
| --- | --- | --- | --- | --- | --- |
| **Study 1 (S1)** | November 20 | 273 students | 70 participants<br>(25% of cohort) | 62 questions | 35-40 minutes |
| **Study 2 (S2)** | April 21 | 273 students | 34 participants<br>(12% of cohort) | 70 questions | 35-40 minutes |
| **Study 3 (S3)** | March 22 | 311 students | 130 participants<br>(42% of cohort) | 34 questions | 20 minutes |
| **Study 4 (S4)** | May 23 | 363 students | 83 participants<br>(23% of cohort) | 38 questions | 20-25 minutes |

**Explanations:** Dates survey conducted, survey lengths and estimated time students required to complete the anonymous JISC online survey. All students enrolled in a bioscience undergraduate degree at Edge Hill University were eligible to participate. The cohort size is the total number of students at the time of the study eligible to participate.

### Quantitative and qualitative data, data interpretation

We conducted four surveys based on a convergent mixed-method approach with all surveys entailing quantitative (single/multiple-choice questions) and qualitative questions (open answer questions) [16]. Upon completion of study, all raw data, each survey layout, and all questions were downloaded from the JISC online survey server and saved on a secure university cloud-based server. Qualitative and quantitative data was collected and analysed separately and combined for *joint display* of results [17]. Quantitative data from all four studies was manually analysed using Microsoft Excel while taking into consideration the grouping, as shown in **table 2**. Each variable/question was analysed independently. Due to the small sample size per group and study, data was not statistically analysed and is presented in raw format, albeit in percentages, allowing easier comparison across all groups, cohorts, and surveys.

**Table 2:** Learn!Bio study: Grouping of Participants.

| **Study year**<br>**(level)** | **Study 1**<br>**(Nov 20)** | **Study 2**<br>**(Apr 21)** | **Study 3**<br>**(Mar 22)** | **Study 4**<br>**(Apr 23)** |
| --- | --- | --- | --- | --- |
| Total number of participants | 70 (100.0%) | 34 (100.0%) | 130 (100.0 %) | 83 (100.0 %) |
| **Year 1 (level 4 – L4)** | 26 (37.1%) | 16 (47.1 %) | 30 (23.1 %) | 27 (32.5 %) |
| **Year 2 (level 5 – L5)** | 28 (40.0 %): | 11 (32.4 %): | 59 (45.4 %) | 41 (49.4 %) |
| **Year 3 (level 6 – L6)** | 16 (22.9%) | 7 (20.6 %) | 41 (31.5 %) | 15 (18.1 %) |

**Explanations:** Studies 1-4 with total numbers of participants per study and grouping into years 1-3 (levels 4-6) for each study. Students' perceptions of learning at the biology department during the Covid-19 pandemic and the following years have been analysed cross-sectional and long-term while comparing overall feedback from participants of classes of 2018-2021 (blue), 2019-2022 (yellow), 2020-2023 (green), 2021-2024 (red), and 2022-2025 (grey). Years 1-3 refer to the UK National Qualification Framework (NQF) levels L4-L6.

Free text/ open questions were reviewed predominantly for internal use to improve teaching and learning at our department and quotes displayed in this manuscript are examples only: Most free-text answer options allowed participants to expand their feedback on topics raised in previously answered quantitative questions, enabled data validation, and provided keywords for coding [18]. All free text answers were analysed individually and across all year with Excel for Microsoft 365 enterprise, version 2402: Numbers of submissions (total, per cohort) were recorded, and each submission was indexed for later identification and publication [19]. Preliminary coding was completed on keywords which arose from the quantitative lead question followed by a detailed coding based on prompts provided by emerging themes (completed by KME).

### Ethical approval

Ethical approval for this study was confirmed by the *Faculty of Arts and Science Research Ethics Committee* (approval reference *FREC/1920/034*) with permission to survey undergraduate students granted by the students’ gatekeeper, Prof Paul Ashton (head of the biology department). We asked students about their place of living, their study programme, if they have care responsibilities (and in Study S1 also about number and age of children they care for), which could lead to indirect identification of participants [20], if combined with other personal data. For this reason, data on gender, sex, age, and origin have not been collected, as per FREC guidance on staff-led studies with student data.

All JISC online surveys (S1-S4) in this study entailed a detailed introduction to the study rationale outlining any ethical dilemmas and how these would be addressed, and a downloadable participant information sheet. Participants were reminded that the survey was completely anonymous, and withdrawal was not possible once the survey had been submitted. On the next online survey page, students were then asked to consent to the JISC online survey by ticking the relevant answers: *I consent to participate in this anonymous survey* - or -*I do not consent to participate in this survey*. If consent was declined, access to the online survey was terminated.

### Survey questionnaires

All survey questionnaires can be found in the **supplements S1**.

## Results

### 1. Learning during the pandemic & adjusting to a new challenging situation

On September 22^nd^, 2020, the UK government announced a new work-from-home policy and introduced a second national lockdown on November 5^th^,2020, followed by student-specific pre-Christmas travel window, enabling students to return to family and friends for the festivities. Covid-19 and frequent regulatory and legal updates resulted in short-notice timetable changes to students returning to academic year 2020-2021 [21–23].

Consequently, students in this study were taught in a blended approach from the start of the 2020/21 academic year with students attending on campus once a week for face-to-face sessions and practical activities, while being taught remotely all other days/sessions. Sessions on campus have been streamed live to enable an all-inclusive learning environment for those unable to attend on campus (shielding, travel restrictions).

This study captured students overall learning experiences after several months of restricted blended learning (November 2020), and then again in March 2022 and April 2023 as detailed in **table 3**. Students were also invited to provide free-text feedback on their overall learning experiences. A first-year biology student suggested this summary on their overall learning experience at the biology department, captured in Study S1 (November 2020):

> *“As someone who has not been on campus due to Covid-19, it has been difficult to keep track of everything and maintain a routine. However, the tutors have been excellent and have worked very hard to ensure things are running smoothly” (L4/9452/5/S1).*

This is in contrast to a comment made by a third-year genetics student in the same study, highlighting the technical and IT hurdles many students encountered whilst learning remotely:

> *“Poor internet connection meant that my mum, brother, and I couldn’t work from home effectively at the same time” [….] (L6/9140/13/S1)*.

**Table 3:** Overall learning experience. Results shown in percentages.

| “Overall learning experience” | Study 1 (Nov 20) |  |  | Study 3 (Mar 22) |  |  | Study 4 (Apr 23) |  |  |
| --- | --- | --- | --- | --- | --- | --- | --- | --- | --- |
|  | L4<br>(26/26) | L5<br>(28/28) | L6<br>(16/16) | L4<br>(30/30) | L5<br>(59/59) | L6<br>(41/41) | L4<br>(27/27) | L5<br>(41/41) | L6<br>(15/15) |
| Excellent | 11.5 | 3.6 | 25.0 | 13.3 | 10.2 | 12.2 | 3.4 | 10.0 | 21.4 |
| Very Good | 46.2 | 25.0 | 6.3 | 43.3 | 42.4 | 31.7 | 37.9 | 15.0 | 42.9 |
| Good | 26.9 | 32.1 | 56.3 | 33.3 | 40.7 | 46.3 | 34.5 | 40.0 | 14.3 |
| Average | 3.8 | 28.6 | 12.5 | 0.0 | 6.8 | 4.9 | 24.1 | 25.0 | 14.3 |
| Poor | 3.8 | 7.1 | 0.0 | 6.7 | 0.0 | 2.4 | 0.0 | 7.5 | 7.1 |
| Very Poor | 0.0 | 3.6 | 0.0 | 3.3 | 0.0 | 2.4 | 0.0 | 2.5 | 0.0 |
| Prefer not to say | 0.0 | 0.0 | 0.0 | 3.3 | 0.0 | 2.4 | 0.0 | 0.0 | 0.0 |
| Overall satisfaction<br>(Excellent – Good) | 84.6 | 60.7 | 87.7 | 90.0 | 93.2 | 90.2 | 75.9 | 85.5 | 78.6 |

**Explanation:** Overall learning experience at the biology department 2020-2023 as experienced by students throughout the Covid-19 pandemic and in follow-up years. Numbers in brackets indicate the total participants followed by total answers provided. Overall satisfaction is the sum of results provided for rubrics “excellent”, “very good”, and “good”. Single-answer question. Results displayed in percentages.

Students and staff worked from home during the third [complete] national lockdown from January 6^th^ to March 7^th^, 2021, which also included the semester 1 exam period. Exams were conducted online as time-limited assessments (TLAs). Staff and year 2 students returned to campus for a (laboratory/fieldwork) research week on March 8^th^, 2021, while all other students continued to work from home until March 15^th^, 2021, when teaching resumed in a blended fashion as described above.

Study S2 was conducted in April 2021, a few weeks after students returned from the third national lockdown (January 6 to March 7^th^) to blended, but still restricted on-campus learning modalities. **Table 4** depicts students’ viewpoints on the impact of the 3^rd^ national lockdown, and the consequential remote-only teaching on their learning.

**Table 4:** Effects of the third national lockdown on learning for bioscience students. Results displayed in percentages.

| “To which extent was your learning affected by the national lockdown in spring 2021” | Study 2 (Apr 21) |  |  |
| --- | --- | --- | --- |
|  | L4 (16/16) | L5 (11/11) | L6 (7/7) |
| Absolutely | 25.0 | 72.7 | 28.6 |
| Partly | 37.5 | 18.2 | 6.3 |
| depends on session content | 37.5 | 9.1 | 12.5 |
| depends on module | 0.0 | 0.0 | 12.5 |
| not at all | 0 | 0 | 0 |

**Explanations:** Students' perceptions of the effects of the UK's 3rd national lockdown (January to March 2021) on their learning at a Higher Education Institution (university). Results from the Learn!Bio study 2 in April 2021. Numbers in brackets indicate the total participants followed by total answers provided. This is a single-answer question. *Results displayed in percentages.*

In April 2021 this study enquired about changes to students’ accommodation provisions as a direct consequence of the Covid-19 pandemic, including campus closures, national lockdowns, or changes in their finances or personal circumstances. **Table 5** outlines changes in accommodation(s) for bioscience students at Edge Hill University in academic year 2020-2021.

Students’ living arrangements in November 2020 are captured in **table S2** in the supplements.

**Table 5:** Covid-19 pandemic and living arrangements for students. Results displayed in percentages.

| “Taken together, how often did you change accommodation this academic year? Please count only | Study 2 (Apr 21) |  |  |
| --- | --- | --- | --- |
|  | L4 (16/16) | L5 (11/11) | L6 (7/7) |

| *study-related changes of accommodation. “* |  |  |  |
| --- | --- | --- | --- |
| Never (0x) | 56.3 | 45.5 | 14.3 |
| Once (1x) | 18.8 | 0.0 | 42.9 |
| Twice (2x) | 25.0 | 36.4 | 28.6 |
| Three times (3x) | 0.0 | 9.1 | 14.3 |
| Four times (4x) | 0.0 | 0.0 | 0.0 |
| More than four times (> 4x) | 0.0 | 9.1 | 0.0 |

**Explanation:** Changes in accommodation as a direct effect of the pandemic in academic year 2020-2021, as reported by bioscience students at the biology department. Numbers in brackets indicate the total participants followed by total answers provided. This is a single-answer question. *Results displayed in percentages.*

Students shared their preferred location for attending remote and blended learning sessions during the autumn 2020 semester, while teaching at most English higher education institutions was restricted to an online or blended teaching manner with very limited access to on-campus facilities and laboratories. At this time bioscience students at our department studied in a remote fashion on 4 out of 5 days and attended on campus on one day per week with different year groups attending on different days to limit the spread of the Sars-Cov2 virus. **Table 6** displays the preferred workplaces for students attending remote teaching in November 2020.

**Table 6:**
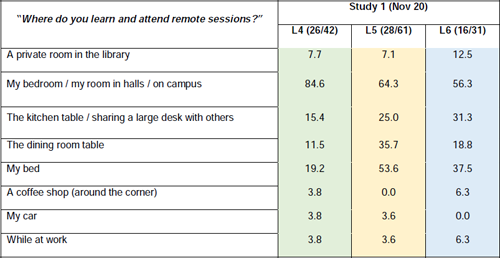
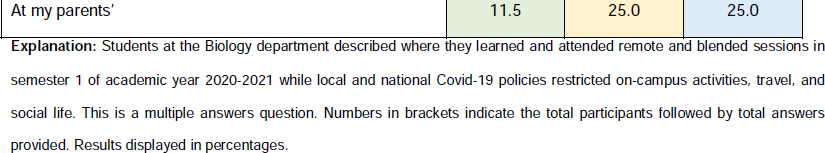
Working at home – students’ workplaces. Results displayed in percentages.

| *“Where do you learn and attend remote sessions?”* | **Study 1 (Nov 20)** |  |  |
| --- | --- | --- | --- |
|  | **L4 (26/42)** | **L5 (28/61)** | **L6 (16/31)** |
| A private room in the library | 7.7 | 7.1 | 12.5 |
| My bedroom / my room in halls / on campus | 84.6 | 64.3 | 56.3 |
| The kitchen table / sharing a large desk with others | 15.4 | 25.0 | 31.3 |
| The dining room table | 11.5 | 35.7 | 18.8 |
| My bed | 19.2 | 53.6 | 37.5 |
| A coffee shop (around the corner) | 3.8 | 0.0 | 6.3 |
| My car | 3.8 | 3.6 | 0.0 |
| While at work | 3.8 | 3.6 | 6.3 |
| At my parents' | 11.5 | 25.0 | 25.0 |

**Explanation:** Students at the Biology department described where they learned and attended remote and blended sessions in semester 1 of academic year 2020-2021 while local and national Covid-19 policies restricted on-campus activities, travel, and social life. This is a multiple answers question. Numbers in brackets indicate the total participants followed by total answers provided. Results displayed in percentages.

Students provided critical feedback on recorded online learning content intended for independent asynchronous study, predominantly made available prior to a scheduled class (**table 7**). Recorded content was made available on the LMS (Learning Management System, Blackboard™) enabled access to all students with internet/IT access.

**Table 7:** Remote learning: Preferences on recorded content. Results displayed in percentages.

| “Which format and style of recorded content do you prefer?” | Study 1 (Nov 20) |  |  |
| --- | --- | --- | --- |
|  | L4 (26/72) | L5 (28/58) | L6 (16/33) |
| Pre-recorded content/presentations are generally well made | 61.5 | 60.7 | 68.8 |
| Pre-recorded content/presentations are often too long (longer than 20 minutes) | 42.3 | 17.9 | 6.3 |
| I struggle with time management if presentations are too long | 38.5 | 21.4 | 12.5 |
| I am ok with longer pre-recorded content/presentations as long as I can stop/start to my needs | 50.0 | 46.4 | 18.8 |
| I am so happy we can now study (more) in our own time - this helps me with my own personal time management | 34.6 | 39.3 | 43.8 |
| Many pre-recorded content/presentations are of poor quality | 7.7 | 0.0 | 0.0 |
| I think pre-recorded content/presentations should not be longer than 5-10 minutes, followed by a quiz | 15.4 | 7.1 | 18.8 |
| I would prefer there is more advice on time-management on when and how to study pre-recorded material | 26.9 | 14.3 | 37.5 |

**Explanation:** Students at the Biology department described their preferences for recorded content made available on the LMS for asynchronous learning prior to a scheduled session. This is a multiple answers question. Numbers in brackets indicate the total participants followed by total answers provided. Results displayed in percentages.

Participants provided 31 free-text responses (15, 11, and 5 responses at L4, L5, and L6, respectively) detailing their views on recorded content in Study S1 (November 2020): Participants preferred recordings which provided closed captures, and showed a clear structure (table of content, slide-by-slide structure). Participants voiced a preference for recordings and PowerPoint slides made available separately, enabling students to *read along* and make annotations while listening to the recordings.

The national and local pandemic policies and regulations in academic years 2019/20 and 2020/21 imposed restrictions on lab-based and field studies. Working at home and attending live online, blended, and recorded sessions from home was an entirely new experience for most of our students during the pandemic. The learning from home experience added unexpected challenges to students’ education and learning, including technical, IT and connectivity issues and persisting struggles with personal well-being, anxiety, and depression, as summarised in **table 8**.

**Table 8:**
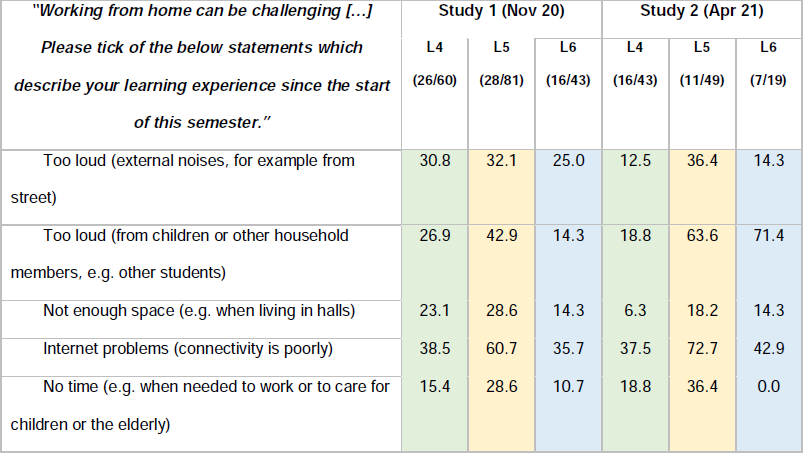
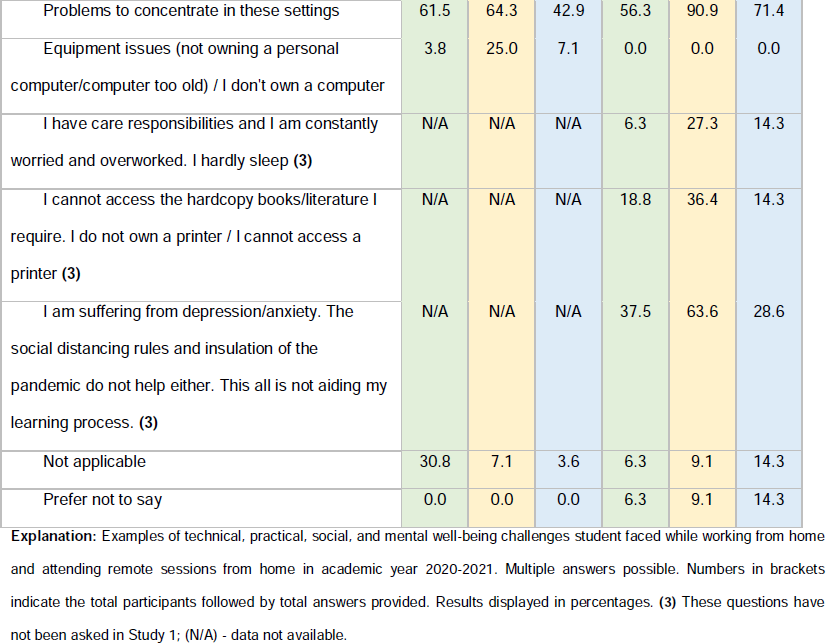
Challenges and hurdles experienced by bioscience students while working from home. Results displayed in percentages.

| ***"Working from home can be challenging [...]***<br>***Please tick of the below statements which***<br>***describe your learning experience since the start***<br>***of this semester."*** | Study 1 (Nov 20) |  |  | Study 2 (Apr 21) |  |  |
| --- | --- | --- | --- | --- | --- | --- |
|  | L4<br>(26/60) | L5<br>(28/81) | L6<br>(16/43) | L4<br>(16/43) | L5<br>(11/49) | L6<br>(7/19) |
| Too loud (external noises, for example from street) | 30.8 | 32.1 | 25.0 | 12.5 | 36.4 | 14.3 |
| Too loud (from children or other household members, e.g. other students) | 26.9 | 42.9 | 14.3 | 18.8 | 63.6 | 71.4 |
| Not enough space (e.g. when living in halls) | 23.1 | 28.6 | 14.3 | 6.3 | 18.2 | 14.3 |
| Internet problems (connectivity is poorly) | 38.5 | 60.7 | 35.7 | 37.5 | 72.7 | 42.9 |
| No time (e.g. when needed to work or to care for children or the elderly) | 15.4 | 28.6 | 10.7 | 18.8 | 36.4 | 0.0 |
| Problems to concentrate in these settings | 61.5 | 64.3 | 42.9 | 56.3 | 90.9 | 71.4 |
| Equipment issues (not owning a personal computer/computer too old) / I don't own a computer | 3.8 | 25.0 | 7.1 | 0.0 | 0.0 | 0.0 |
| I have care responsibilities and I am constantly worried and overworked. I hardly sleep **(3)** | N/A | N/A | N/A | 6.3 | 27.3 | 14.3 |
| I cannot access the hardcopy books/literature I require. I do not own a printer / I cannot access a printer **(3)** | N/A | N/A | N/A | 18.8 | 36.4 | 14.3 |
| I am suffering from depression/anxiety. The social distancing rules and insulation of the pandemic do not help either. This all is not aiding my learning process. **(3)** | N/A | N/A | N/A | 37.5 | 63.6 | 28.6 |
| Not applicable | 30.8 | 7.1 | 3.6 | 6.3 | 9.1 | 14.3 |
| Prefer not to say | 0.0 | 0.0 | 0.0 | 6.3 | 9.1 | 14.3 |

**Explanation:** Examples of technical, practical, social, and mental well-being challenges student faced while working from home and attending remote sessions from home in academic year 2020-2021. Multiple answers possible. Numbers in brackets indicate the total participants followed by total answers provided. Results displayed in percentages. **(3)** These questions have not been asked in Study 1; (N/A) - data not available.

Feedback from staff facilitating online and blended learning experiences initiated a question on hurdles to microphone and camera use in the remote learning in the April 2021 (Study S2). Students’ responses are shown in **figure S3** in the supplements.

With only limited on-campus sessions available for most of academic years 2019/20 and 2020/21 many students had reported a lack of structure and the wish for additional guidance for their remote learning routines, represented in a statement by a year 1 plant science student captured in Study S1 (November 2020):

> *“[….] Clear, concise [work] instructions for everything” (L4/3902/62/S1).*

A fellow first year student, studying towards a bachelor’s degree in biomedical science, summarised their feelings about new and unknown staff and the challenges of all-remote modules in a statement captured in November 2020:

> *“I feel that having set modules which are taught exclusively online is negatively affecting my studies, I find it harder to understand and harder to ask for help having never met those [who] are teaching me, maybe at least a few sessions in each module could be taught face to face to assist [my] learning” (L4/6421/62)*.

In November 2020 we asked students about their newly adopted learning, planning, and assessment preparation strategies as a consequence of the pandemic and the need to work from home (multiple-answer question, **table 9**).

**Table 9:**
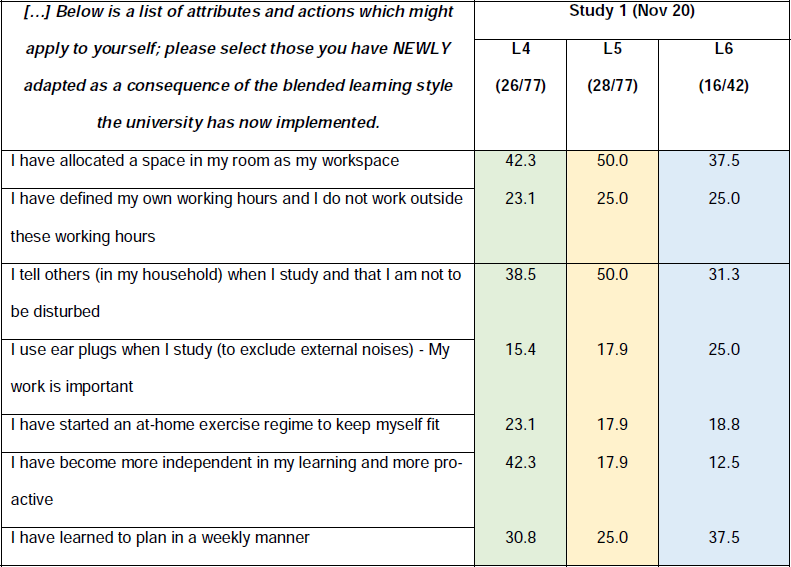
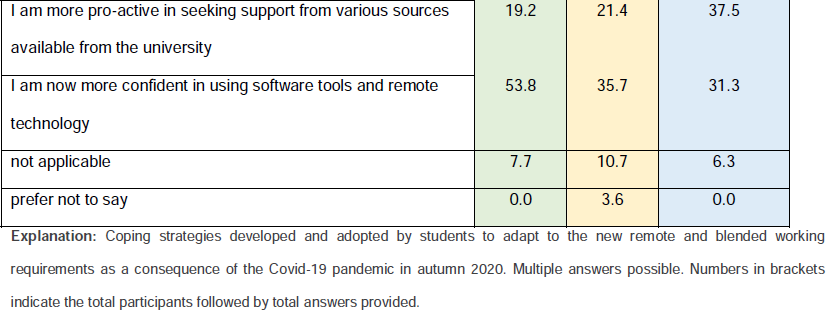
Skills development and coping strategies as a consequence of the pandemic. Results displayed in percentages.

| [...] Below is a list of attributes and actions which might apply to yourself; please select those you have **NEWLY** adapted as a consequence of the blended learning style the university has now implemented. | Study 1 (Nov 20) |  |  |
| --- | --- | --- | --- |
|  | L4<br>(26/77) | L5<br>(28/77) | L6<br>(16/42) |
| I have allocated a space in my room as my workspace | 42.3 | 50.0 | 37.5 |
| I have defined my own working hours and I do not work outside these working hours | 23.1 | 25.0 | 25.0 |
| I tell others (in my household) when I study and that I am not to be disturbed | 38.5 | 50.0 | 31.3 |
| I use ear plugs when I study (to exclude external noises) - My work is important | 15.4 | 17.9 | 25.0 |
| I have started an at-home exercise regime to keep myself fit | 23.1 | 17.9 | 18.8 |
| I have become more independent in my learning and more proactive | 42.3 | 17.9 | 12.5 |
| I have learned to plan in a weekly manner | 30.8 | 25.0 | 37.5 |
| I am more pro-active in seeking support from various sources available from the university | 19.2 | 21.4 | 37.5 |
| I am now more confident in using software tools and remote technology | 53.8 | 35.7 | 31.3 |
| not applicable | 7.7 | 10.7 | 6.3 |
| prefer not to say | 0.0 | 3.6 | 0.0 |

**Explanation:** Coping strategies developed and adopted by students to adapt to the new remote and blended working requirements as a consequence of the Covid-19 pandemic in autumn 2020. Multiple answers possible. Numbers in brackets indicate the total participants followed by total answers provided.

Survey participants voiced their opinion on how the department could assist students in their eLearning and might offer coping strategies in 38 free text responses (L4: 14, L5: 16, L6: 8) with 42.1% (16/38) responders expressing a wish for more non-modular and fun activities offered online, 34.2% (13/38) against, and 26.3% (10/38) outlining other/personal statements and feedback; for example a year 1 genetics student expressing their wish for more extra-modular activities:

> “*Both during lectures and within separate bioscience group chats we only talk about academic topics and do not know each other well (especially those from other teaching groups). I find most of my social interaction/people I know best through societies, so maybe having optional fun sessions could help students interact with each other more” (L4/0711/57/S1)*.

A year 2 biology student living in private accommodation in the nearby town of Ormskirk expressed a different opinion:

> “*I live alone and the only time I talk to anyone since I’ve come back to Ormskirk is for a few minutes before I go into the [X] lecture on Thursday. I spend 24 hours of most days in a 4m x 4m room alone. I don’t really think extra sessions would make much of a difference for me personally” (L5/0378/57/S1)*.

For any on-campus teaching in semester 1 in academic year 2020/21, students were allocated into small groups (“bubbles”) allowing the required two meter distanced spacing during classroom teaching. Each student received their own set of equipment for any lab experiment. In the November 2020 survey (Study S1) students have been asked if they felt safe with the offered on-campus teaching arrangements; 88.5% of all L4 students (28 participants providing 28 responses) fully/to some extent, agreed that it was safe to be in a classroom, as well as 85.7% of all L5 students (26/26) and 62.7/% of L6 students (16/16).

Students acknowledged the extra workload that technical staff were experiencing as 92.3%, 85.8%, and 68.8% students L4-L6 fully or to some extent agreed that “*technical staff do an amazing job preparing each session*”. Students in levels L4-L6 agreed fully or to some extent with 96.2%, 96.4%, and 75%, respectively that “*all staff do they best they can, given the circumstances*”.

### 2. Wellbeing & resilience

The advent of the Covid-19 pandemic and the restrictions implemented on national and local levels resulted in extensive uncertainty amongst our students. The third national lockdown in spring 2021 (Jan 6^th^ – March 7^th^) resulted in an exponential decrease in students’ wellbeing (Study S2), with 43.7% of level 4 (L4) students, 72.5% of level 5 (L5), and 42.9% of level 6 (L6) students reporting of “poor” or “very poor” mental health (**figure 1**).

**Figure 1:**
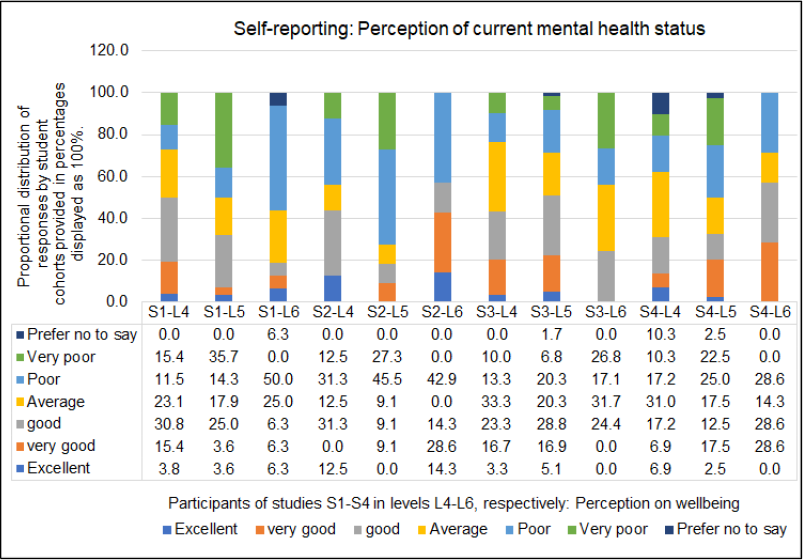
Participants perception of their own current wellbeing from November 2020 (Study 1) to May 2023 (Study 4). Students have been surveyed in November 2020 (S1-L4, S1-L5, S1-L6), April 2021 (S2-S4, S2-L5, S2-L6), March 2022 (S3-L4, S3-L4, S3-L6) and May 2023 (S4-L4, S4-L5, S4-L6). For cohort sizes and student grouping see table 1. Single-answer question. Data shown and displayed in percentages.

In all follow-up surveys 2021-2023 students reflected on changes in their mental health as compared to the past semester (**table 10**). All year groups reported a “large” mental health decline from semester 1 to semester 2 in academic year 2020-2021 (study conducted shortly after the third national lockdown). Such a prominent change in wellbeing was not observed again during the lifetime of this study.

**Table 10:** How mental health changed compared to last semester. Results displayed in percentages.

| Change in mental health past semester to this semester | Study 2 (Apr 21) |  |  | Study 3 (Mar 22) |  |  | Study 4 (Apr 23) |  |  |
| --- | --- | --- | --- | --- | --- | --- | --- | --- | --- |
|  | L4 | L5 | L6 | L4 | L5 | L6 | L4 | L5 | L6 |
|  | (16/16) | (11/11) | (7/7) | (30/30) | (59/59) | (41/41) | (27/27) | (41/41) | (15/15) |
| large improvement | 12.5 | 9.1 | 57.1 | 10.0 | 22.0 | 14.6 | 6.9 | 12.5 | 0.0 |
| small improvement | 6.3 | 9.1 | 14.3 | 0.0 | 0.0 | 0.0 | 20.7 | 17.5 | 21.4 |
| unchanged | 31.3 | 9.1 | 0.0 | 30.0 | 30.5 | 12.2 | 20.7 | 15.0 | 42.9 |
| small decline | 6.3 | 9.1 | 14.3 | 26.7 | 10.2 | 24.4 | 27.6 | 30.0 | 35.7 |
| large decline | **25.0** | **45.5** | **28.6** | 3.3 | 1.7 | 2.4 | 13.8 | 17.5 | 0.0 |
| unsure | 31.3 | 36.4 | 14.3 | 33.3 | 37.3 | 48.8 | 3.4 | 2.5 | 0.0 |
| prefer not to say | 6.3 | 9.1 | 14.3 | 0.0 | 0.0 | 0.0 | 6.9 | 5.0 | 0.0 |

**Explanation:** Participants' perception of changes to their mental health in comparison to the previous semester. Single answer question. Numbers in brackets indicate the total participants followed by total answers provided. Results displayed in percentages.

Students reported a range of variables which might have affected their wellbeing and mental health in academic year 2020-2021, most prominently the lack of opportunities for communication with others (students, adults, peers), and missing a community feeling, which was even more evident after the third national lockdown (Jan 6^th^ to March 7^th^, 2021). From spring 2021 onwards, students were more exposed to bereavements caused by the SARS-CoV-2 virus infections and their consequences within close family or friends (**table 11**).

**Table 11:** Consequences of remote and blended learning on participants wellbeing in academic year 2020-2021. Results displayed in percentages.

| ***“Do you believe the new blended learning approach, with less on campus and more remote teaching has an impact on your mental wellbeing? Please tick all statements which apply to your situation.”*** | **Study 1 (Nov 20)** |  |  | **Study 2 (Apr 21)** |  |  |
| --- | --- | --- | --- | --- | --- | --- |
|  | **L4** | **L5** | **L6** | **L4** | **L5** | **L6** |
|  | **(26/59)** | **(28/108)** | **(16/56)** | **(16/50)** | **(11/56)** | **(7/19)** |
| I feel lonely /more lonely | 11.5 | 46.4 | 50.0 | 31.3 | 54.5 | 28.6 |
| I miss a community feeling | 38.5 | 42.9 | 43.8 | 50.0 | 54.5 | 42.9 |
| I miss a feeling of belonging | 7.7 | 35.7 | 31.3 | 37.5 | 45.5 | 71.4 |
| I miss talking to people | 38.5 | 64.3 | 62.5 | 50.0 | 54.5 | 42.9 |
| I miss speaking to adults | 7.7 | 32.1 | 12.5 | 18.8 | 45.5 | 14.3 |
| I miss to relax /chill out with others | 30.8 | 46.4 | 56.3 | 25.0 | 63.6 | 28.6 |
| I feel depressed /anxious | 11.5 | 32.1 | 18.8 | 31.3 | 81.8 | 14.3 |
| I am anxious about the future/my studies | 26.9 | 64.3 | 56.3 | (#) | (#) | (#) |
| I am suffering from severe anxiety I am suffering from a clinically proven depression | (*) | (*) | (*) | 50.0 | 63.6 | 28.6 |
| There have been one or more bereavements affecting me/my family/my close friends | (*) | (*) | (*) | 18.8 | 54.5 | 14.3 |

**Explanation:** Multiple answers possible. Numbers in brackets indicate the total participants followed by total answers provided.
Results displayed in percentages. (#) question not asked in Study 2 (April 21), (\*) question not asked in Study 1 (November 20).

During times of remote or blended learning students were supported by their respective year coordinator, personal tutors, and module leaders at the biology department, who all reached out to students on a regular basis. Frequent changes in national and local Covid-19 policies and regulations resulted in ad hoc timetable adjustments with often significant consequences for students’ learning experiences. In the April 2021 Study S2 feedback was sought on participants’ achievements on building-up resilience in these challenging times and how students perceived their own resilience at this stage (**table 12**). Additional supportive information on resilience development can be found in **supplemental table S4.**

**Table 12:**
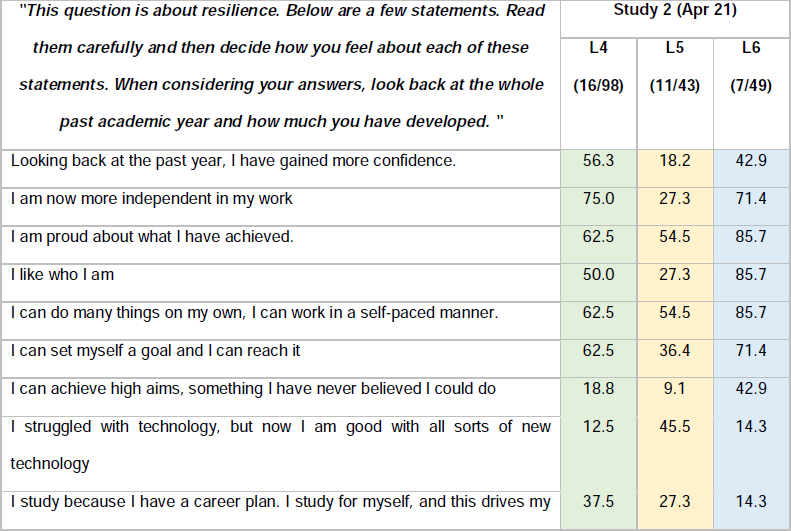
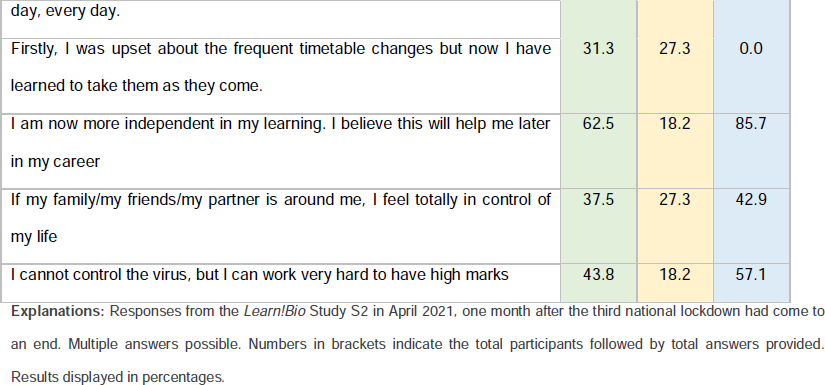
Participants pathway to resilience at the height of the Covid-19 pandemic. Results displayed in percentages.

### 3. Consequences of the pandemic & students’ expectations from providers

We conducted two post-pandemic follow-up surveys shedding light on long-term consequences for higher education bioscience students on learning experiences and wellbeing because of the Covid-19 pandemic. The first post-pandemic survey took place in March 2022 (Study S3) followed by a final survey in April 2023 (Study S4). Participants provided feedback on the (long-term) effects of (pandemic) online and blended teaching on their current learning progression (**table 13**).

**Table 13:** Long-term effects of remote learning on current bioscience students. Results displayed in percentages.

| “Do you feel the past remote and blended learning during the pandemic affects your current learning?” | Study 3 (Mar 22) |  |  |
| --- | --- | --- | --- |
|  | L4<br>(30/30) | L5<br>(59/59) | L6<br>(41/41) |
| Absolutely | 16.7 | 35.6 | 41.5 |
| Partly | 20.0 | 23.7 | 19.5 |
| Depends on session content | 26.7 | 16.9 | 17.1 |
| Depends on module | 13.3 | 11.9 | 12.2 |
| Not at all | 23.3 | 10.2 | 9.8 |

**Explanation:** Consequences of online and remote teaching during the Covid-19 pandemic on current learning progression as perceived by students at the biology department at Edge Hill University captured at the post-pandemic follow-up survey in 2022 (Study S3). Single answer questions. Numbers in brackets indicate the total participants followed by total answers provided. Results displayed in percentages.

As a result of the Study 3 preliminary data analysis (**table 13**) and conversations with students we extended our questionnaire for the final survey in April 2023 allowing additional insight on participants’ pre-university learning restrictions during the Covid-19 pandemic (**table 14**, **table 15**).

**Table 14:** Effects of students pre-University learning on current university progression. Results displayed in percentages.

| ***"Covid-19 and the pandemic: Has your pre-Uni and early-university learning had an impact on your current studies?"*** | **Study 4 (Apr 23)** |  |  |
| --- | --- | --- | --- |
|  | **L4**<br>**(27/27)** | **L5**<br>**(41/41)** | **L6**<br>**(15/15)** |
| Absolutely | 27.6 | 40.0 | 28.6 |
| Partially | 27.6 | 40.0 | 28.6 |
| Not at all | 34.5 | 15.0 | 28.6 |
| Unsure | 3.4 | 2.5 | 14.3 |
| Prefer not to say | 6.9 | 2.5 | 0.0 |

**Explanation:** Single answer questions. Numbers in brackets indicate the total participants followed by total answers provided. Results displayed in percentages.

**Table 15:**
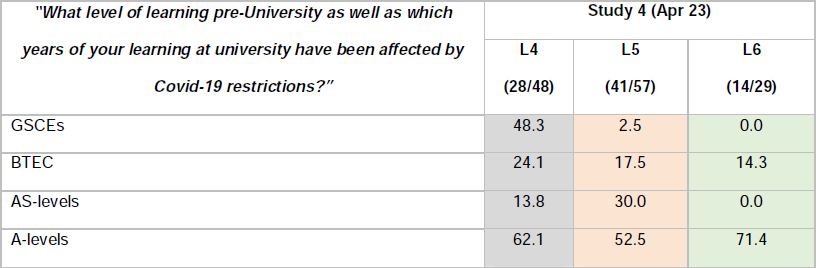
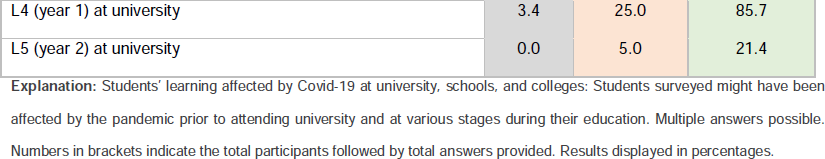
Level of learning affected by Covid-19. Results displayed in percentages.

Throughout the pandemic, students acquired various new IT and computer skills. In our follow-up surveys in 2022 and 2023, respectively, we sought feedback from participants how they felt about their IT literacy and technology advancements (**table 16**).

**Table 16:** IT technology & the post-pandemic generation of HE learners. Results displayed in percentages.

| Do you feel you are now more technology advanced? | Study 3 (Mar 22) |  |  | Study 4 (Apr 23) |  |  |
| --- | --- | --- | --- | --- | --- | --- |
|  | L4 (30/30) | L5 (59/59) | L6 (41/41) | L4 (27/27) | L5 (41/41) | L6 (15/15) |
| Yes | 86.7 | 81.4 | 75.6 | 85.7 | 87.8 | 92.9 |
| No | 10.0 | 16.9 | 24.4 | 10.7 | 0.0 | 0.0 |
| Unsure | n.a. | n.a. | n.a. | 7.1 | 9.8 | 7.1 |
| Prefer not to say | 3.3 | 1.7 | 0.0 | n.a. | n.a. | n.a. |

**Explanation:** Technology advanced learners. Participants reflection on their ability to use computers, and online technology as compared to before the pandemic with remote and blended learning requiring them to embrace these new tools and technology. (n.a.) - not asked. Single answer questions. Numbers in brackets indicate the total participants followed by total answers provided. Results displayed in percentages.

Students in this study returned to pre-pandemic, on-campus learning setting from September 2021, as requested by the Office for Students (OfS) and enabled by national Covid-19 regulations [24]. With digital teaching technology now widely available enabling colleagues to deliver blended and online sessions, students’ perceptions on future teaching practices were captured in Study 3 (2022) and Study 4 (2023). Students express a clear preference for a combination on blended and face-to-face teaching but rated purely online teaching less favourably (**table 17**). Preferences have not changed from 2022 to 2023.

**Table 17:** Students preferences on teaching post-pandemic: Results displayed in percentages.

| *“Please indicate your preferred teaching and learning style at university”* | Study 3 (Mar 22) |  |  | Study 4 (Apr 23) |  |  |
| --- | --- | --- | --- | --- | --- | --- |
|  | L4<br>(30/30) | L5<br>(59/59) | L6<br>(41/41) | L4<br>(28/28) | L5<br>(41/41) | L6<br>(14/14) |
| Online learning (off campus) | 0.0 | 1.7 | 0.0 | 10.3 | 2.5 | 0.0 |
| **blended learning** | **56.7** | **49.2** | **41.5** | **44.8** | **42.5** | **50.0** |
| **face-to-face (on campus)** | **36.7** | **47.5** | **58.5** | **44.8** | **55.0** | **50.0** |
| no preference | 6.7 | 1.7 | 0.0 | 0.0 | 0.0 | 0.0 |

**Explanation:** Bioscience students' preferences for online, blended, and face-to-face (on campus) teaching as identified in the S3 and S4 surveys, respectively. Single answer questions. Numbers in brackets indicate the total participants followed by total answers provided. Results displayed in percentages.

Students were surveyed about which of the non-LMS embedded learning tools and software introduced for remote and blended learning during the Covid-19 pandemic, that the department should continue to use in its teaching delivery. Students from all cohorts mentioned YouTube (54.7%, range 48.8-65.5%), Kahoot! (49.4%, range 39.9-52.5%), LinkedIn Learning (8.9%, range 3.3-26.8%), Socrative (11.5%, range 3.3-20.3%), Padlet (10.1%, range 0.0-29.3%), and Clinical Key Students (12.3%, range 3.3-26.8%). Not all students in all year groups have had previous experience with all mentioned online learning tools.

In April 2023 (Study S3) participants provided insights to which extend the consequences of the pandemics still affected their [current] social life (**table 18**).

**Table 18:** Social life consequences of the Covid-19 pandemic. Results displayed in percentages.

| *“Has the pandemic affected your social life?”* | Study 4 (Apr 23) |  |  |
| --- | --- | --- | --- |
|  | L4<br>(30/30) | L5<br>(59/59) | L6<br>(41/41) |
| Yes, by a large amount | 31.0 | 30.0 | 42.9 |
| yes, by a small amount | 27.6 | 30.0 | 57.1 |
| no change | 37.9 | 27.5 | 0.0 |
| unsure | 3.4 | 12.5 | 0.0 |

**Explanation:** Long-term effects of the Covid-19 pandemic on bioscience students' social life surveyed in April 2023. Single answer questions. Numbers indicate the total participants followed by total answers provided. Results displayed in percentages.

Amongst all participants surveyed in this study at level L4, 56% report about anxiety (disorders) and/or autism spectrum disorders (ASD), as well as 38% at level L5, and 42% at level L6, respectively. A year 2 biotechnology student expressed their struggles with on-campus learning in this statement, captured in the Study S3 (March 2022):

> *“[…] The long drive to uni[versity] affects my ability to learn at in-person lectures - I feel, much like last year, I would learn a lot more sat at home in my own environment. […] As someone with autism it’s difficult to form new routines and sometimes, as silly as this sounds, my entire day can be ruined from just one thing like someone sitting in a seat I usually sit in which is why it’s important to ensure that online learning is still an option when it comes to lectures as, with some of us, the commute can seriously impact our ability to learn” (L5/5418/26a/S3)*.

In the post-pandemic follow-up surveys S3 and S4, conducted 2022 and 2023, respectively, students’ perception on possible social and personal factors which might affect their current learning and hindrances to return to university were captured (**table 19**).

**Table 19:** Personal and social attributes and how these inform student’s perception of post-pandemic teaching and learning. Results displayed in percentages.

| *"Reflect on your approach to teaching and learning this academic year."* | Study 3 (Mar 22) |  |  | Study 4 (Apr 23) |  |  |
| --- | --- | --- | --- | --- | --- | --- |
|  | L4 | L5 | L6 | L4 | L5 | L6 |

|  | (30/79) | (59/178) | (41/119) | (27/44) | (41/63) | (15/24) |
| --- | --- | --- | --- | --- | --- | --- |
| I am excited to be getting back to pre-pandemic learning | 50.0 | 54.2 | 53.7 | 3.7 | 24.4 | 40.0 |
| I am anxious about being in contact with large number of people during a pandemic | 30.0 | 27.1 | 14.6 | 14.8 | 12.2 | 6.7 |
| I am I excited to meet my class/roommates in person | 36.7 | 39.0 | 56.1 | 18.5 | 9.8 | 13.3 |
| I am anxious about the lack of social distancing/mask wearing on campus/my commute | 10.0 | 15.3 | 4.9 | 7.4 | 4.9 | 6.7 |
| I watched videos during remote learning to gain practical knowledge, now I feel I am prepared to learn and develop those lab skills. | 10.0 | 13.6 | 4.9 | 7.4 | 7.3 | 6.7 |
| **I am anxious about my lack of practical knowledge from missed in-person teaching** | **30.0** | **52.5** | **46.3** | **25.9** | **24.4** | **26.7** |
| The lockdowns gave me a nice break from the everyday social situations | **53.3** | **64.4** | **61.0** | 14.8 | 22.0 | 20.0 |
| The lockdowns have made me more self-conscious in everyday social situations | 30.0 | 33.9 | 43.9 | 37.0 | 17.1 | 26.7 |
| N/A - There has been no impact on my learning | 13.3 | 1.7 | 4.9 | 18.5 | 26.8 | 0.0 |

**Explanation:** Multiple answers possible. Numbers in brackets indicate the total participants followed by total answers provided.
Results displayed in percentages.

In the 2022 follow-up study (Study S3) students still felt the imminent effects of the pandemic and voiced their estrangement from a pre-pandemic social behaviour, which was less evident in the 2023, Study 4.

We captured our final year undergraduate bioscience students’ career plans and how these have been impacted by the Covid-19 pandemic and its consequences on access to training, job-preparedness, and availability of graduate bioscience roles (**table 20**). During the pandemic we observed that a higher than usual number of students continued with their education (MRes, MSc, PhD), either with us or elsewhere instead of pursuing a paid-for career. A year 3 (L6) ecology & conservation student’s voice from the November 2020 (Study S1) summarises this reasoning:

> “[The] *change of dissertation topic meant I had interest in a different postgraduate course (…), also I feel by doing a master [degree] I avoid the current situation with a lack of jobs (L6/0169/61/S1)*”.

A human biology student in their final year expressed their worries about lack of job vacancies for bioscience graduates in the March 2021 Study S2:

> *“I am applying for full-time jobs within the NHS, Civil Service, local universities, and other public services. I will continue working my part-time job until I secure a full-time position. […] I am concerned about high levels of unemployment at the minute making securing a job very difficult. I have already applied to a large number of jobs that I am sufficiently qualified for and have been rejected by every one of them. I think it is going to be a long process” (L6/2144/69/S2)*.

**Table 20:**
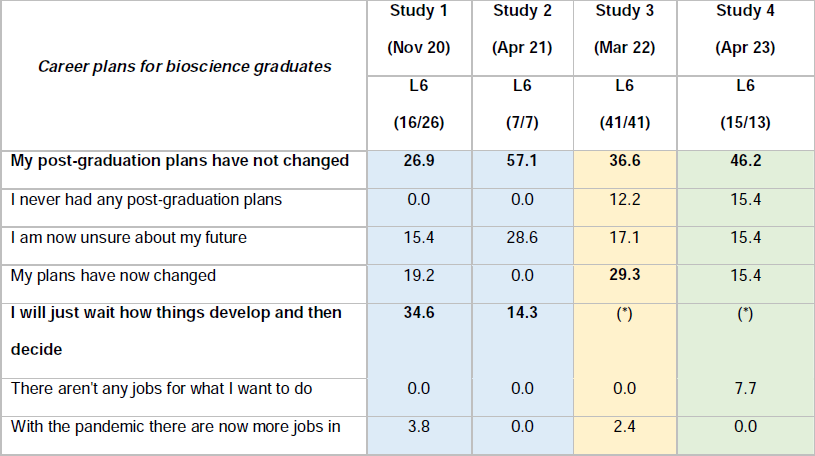
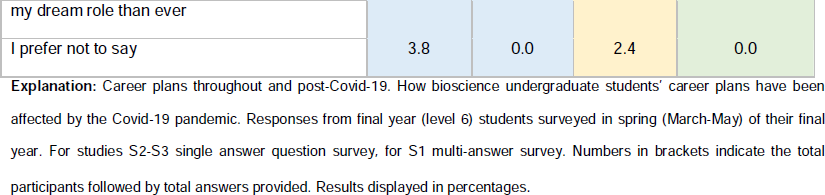
Bioscience students career plans during and after the Covid-19 pandemic. Results displayed in percentages.

## Discussion

### 1. Learning during the pandemic & adjusting to a new challenging situation

This manuscript evaluated students’ perception on studying and completing a bioscience degree during and post the Covid-19 pandemic from November 2020 to March 2023 in a cross-sectional, mixed-method approach by inviting undergraduate students to four independent, anonymous JISC online surveys as part of our departmental *Learn!Bio* study.

We firstly evaluated bioscience students’ perceptions on their learning experiences and how students adjusted to the practical and learning-relevant challenges which had arisen in the wake of the Covid-19 pandemic from November 2020 to spring 2021. All undergraduate students enrolled at the biology department at EHU were eligible to participate. Students were invited via a mass email sent to their student email inbox, which entailed a link to the anonymous JISC online survey. Students received a reminder email two weeks later.

Edge Hill University is a located in West Lancashire with a predominantly White (English, Welsh, Scottish, Northern Irish or British) population (96.9%, 2021 census [25]) population. Asian, Asian British or Asian Welsh citizens and Black, Black British, Black Welsh, Caribbean, or African citizens account for only 1.05% and 0.33% of the local community (2021 census [25]), respectively. Only 4.3% of the local population are non-UK passport holders (Manchester: 19.2%, City of London: 34% [26]). There is a significant internal disparity in income distribution in West Lancashire, which is also reflected in the university’s student cohort, with 15 out of 273 neighbourhoods ranked amongst the 20% most deprived with an income deprivation of 39%. In contrast 11/273 neighbourhoods in West Lancashire are ranked amongst the least deprived in England with an income deprivation of 2.7% [27]. The university attracts a high number of local students who continue to live at home during their studies.

Study satisfaction captured amongst the students in the November 2020 (Study S1, **table 3**) was well above the national average, when compared with data from a national-wide Student Covid Insights Survey (SCIS) collected in October and November 2020 showing a 29% rate of dissatisfaction or severe dissatisfaction amongst surveyed students with their studies (71% satisfaction rate [12]). This contrasts with 88.1% and above of satisfaction amongst students in this study.

Most students in this study felt they have been affected in their learning by the national lockdown and the online-only teaching in spring 2021 (**table 4**). This result is supported by national-wide evidence from the 2021 National Student Survey (NSS) showing that only 47.6% of all respondents have been content with the delivery of their learning and teaching during the Covid pandemic, which was further emphasised by data made available from the Student Covid-19 Insights Survey (SCIS), reporting that 52% of students who had been in higher education during the pandemic felt that Covid-19 had a “significant” or “major impact” on their academic performance [12, 28, 29].

Above national average of participants captured in this manuscript changed accommodation at least once during academic year 2020/21 (**table 5**), with many students moving between their “home” and “study accommodation”. Nation-wide 81% of all students remained in their study-related accommodation in academic year 2020/21 [28].

A high percentage of students in this cohort lived off campus, predominantly with their parents (43%) or with partners/with their own children (18%) during the height of the pandemic, a factor which might has decreased their satisfaction with their studies, as space and IT resources for learning might have been restricted. A recent study by colleagues from the Aston University has shown that students from more deprived households have rated their study experiences less favourable during the pandemic, as these students have more frequently been deprived in one or more dimension [30], and might have lacked access to their own computer, fast internet or a suitable learning environment.

Accessing timetabled eLearning sessions with a lack of technology and/or a quiet space to concentrate was identified as a main hurdle for many participants as captured in November 2020 (Study S1). Access to a personal computer, computer accessories (microphone, camera) and internet connectivity was also a worry for many participants (**table 8**). Our university, as many other HEIs in the UK, provided students in need, with laptops and associated equipment (cameras, headsets etc) for long-term loan, along with internet dongles to enable equal access to eLearning for all students. This resulted in the eradication of unobtainable access to personal computers by March 2021 (**table 8**).

Staff at our university offered a mixture of synchronous and asynchronous teaching approaches with pre-recorded podcasts and presentations made available in preparation for on-campus or remote synchronous session. Most students agreed that remotely accessible learning content was generally well made (**table 7**), a viewpoint also observed by other studies in the UK and globally [30, 31]. Students in year 1 (L4) had only little opportunity to acquire the endurance and attention span to engage with lengthy learning content during their often-interrupted school/college education, which might explain their reluctance to engage with longer (20 minutes-plus) remote learning units [32].

### 2. Wellbeing & resilience

An NHS digital survey analysis conducted on members of the public living in England and published in 2016 concluded that one in five women and one in eight men suffered from a common mental health condition prior to the Covid-19 pandemic [33]. Anxiety and symptoms of anxiety have been more evident in the UK general population before the Covid-19 pandemic (21%) compared to other countries, for instance Belgium (11%), New Zealand (6.1 %), the USA (8.2%) or Sweden (14.2%), as recently reported by The King’s Fund (see reference [34], pages 33 and following). The marked differences in levels of anxiety and related conditions experienced by members of the public in the UK as compared to members of the public in other countries increased during the Covid-19 pandemic, with only 30.8% of the public in the USA expressing symptoms of anxiety, and 22.3% of the public in EU member states (averaged) reporting of anxiety symptoms during these times. This compares to 49.6% of the public in the UK with anxiety symptoms during the pandemic [34, 35].

The UK experienced a full, strict national Covid-19 lockdown from January 6^th^ to March 7^th^, 2021.

Shortly after the third national lockdown, in April 2021, data for the second survey (Study S2) was captured, which showed a marked decrease in students overall wellbeing (**figure 1**). Students at level 5 (class of 2019-2022, shown in yellow), reporting well below the national average on poor mental health as compared to all UK adults. This student cohort’s mental health, captured at this time point was also well below the reported average as reported in the April Student Covid-19 Insights Survey, in which 63% of all students expressed “*slightly or much worse now*” mental health since their started their studies in autumn 2020 [28] This trend in mental health decline amongst university students was also identified in a recent study by Chen and Lucock [36].

Students of the class 2019-2022 (displayed with yellow background) have overall reported more critical feedback and negative impressions in all surveys S1-S3. The class of 2019-2022 experienced Covid-19 restrictions throughout all years of their undergraduate studies, from the spring semester 2020 to summer 2022. This was also evidenced in students’ perception on their mental health changes from autumn 2021 to spring 2022 (**table 10**), with students in the cohort of 2019-2022, showing a more severe decline in their wellbeing compared to students in the class of 2020-2023 cohort (displayed with green background). Nation-wide, students’ mental health experienced a further decline from autumn 2021 to spring 2022, as evidenced in the SCIC March 2022 survey [37].

Recent studies have shown that restrictions to campus life and student services affected those students from disadvantaged socioeconomic backgrounds, with often poor “*networking capital*” [11] and learners with (child) care responsibilities or inadequate IT equipment or internet connectivity, more severely [36, 38]. Interestingly, our data indicates that campus closures and restrictions to libraries, IT rooms and on-campus teaching had a cross-sectional impact on all students in all years, except for the 2020-23 cohort (shown with green background) captured in the November 2020 study (S1), who might have capitalised on their previous (school/college) social network structures for support and wellbeing (**table 11**) during their first term at university.

In the April 2020 survey (S2) we captured students’ perception on the development of new skills and students’ pathway to resilience (**table 12**, supplementary **figure S4**). Feedback from students on their resilience and acquired skills seems to conflict with an overall perception of their current wellbeing at this time (“poor”, “very poor”), as students in years 1 and 3 (levels 4 and 6) provided a positive self-rating for their transition to resilience and have recognised the acquisition of various new soft and IT skills and as wells and job-relevant graduate attributes required for a role in biosciences. Students in year 2 (level 5) have been more critical about their skills development and resilience advancements while these students expressed the most significant decline in mental health across all year groups.

### 3. Consequences of the pandemic & students’ expectations from providers

In two follow-up, post-pandemic surveys this manuscript captured students’ perception on long-term consequences of blended and remote learning on their bioscience education, wellbeing, and social life in academic years 2021-2022 (Study S3) and academic year 2022-2023 (Study S4). Responses show that 76.7%, 88.1%, and 90.2% of all students in years 1, 2, and 3, respectively, indicated that the previously experienced Covid-19 pandemic restrictions are affected their higher education in 2022 (**table 13**). Interestingly this effect continued to impact 51.9%, 56.1%, and 86.7%, of all year 1, 2, and 3 students, respectively one year later (Study S4, **table 14**).

Covid-19 has impacted not only students’ higher education but also their school and college education (**table 15**); with those from the most deprived backgrounds reporting more often that they have fallen behind (42%) than those from the least deprived backgrounds (26%) [39]. Many if not all outreach activities had been cancelled during the pandemic years, affecting career choices and accession to university programmes, with those from deprived areas more affected than others [38]. Our findings are supported by a recent meta-analysis investigating student learning outcome across various countries by De Pietro *et al*. concluding with a significant learning deficit in math and science, as a consequence of pandemic learning, indicating that long-term support in STEM subjects might be required for school and early-stage university students, allowing equal and fair chances for all learners catching up with their learning [40].

In our study, students’ perception on their IT and computer literacy has fulminant increased from November 2020 to May 2023 with 92.9% of the students in the 2020-2023 cohort (green) self-reporting on improved IT skills (up from 12.5%, **table 12 vs table 16**), indicative of a pathway to resilience and an adaption to a new learning environment. This is further evidenced by feedback provided to the question on how future teaching should be mediated (online, blended, campus-based/face-to-face). Students are supportive of both, face-to-face and blended synchronous teaching, with blended/hybrid teaching allowing students to attend live streamed sessions remotely (**table 17**). Our department, as most HEIs in the UK require students to attend physically for all scheduled sessions. In various free text responses students reported about their reasons not to attend on campus, including public transport not available at all/on certain days/impacted by strike actions, care responsibilities, sickness (including mental health and chronic conditions). Many students expressed worries about the significantly increased living costs the impact their travel arrangements to/from campus for “*just this session*”.

The financial aspects of attending university sessions are also captured in the recent the ONS Student Cost of Living Insights Study (SCoLIS) from spring 2023, which suggested that 91% of all students worry/worry very about their finances and that 49% of all students have financial difficulties [41]. A recent study by Salem and colleagues pinpoints to evidence on best learning outcomes with blended learning strategies while critically arguing for highest students’ overall motivation and satisfaction through face-to-face learning modalities [42].

The Covid-19 pandemic has increased the prevalence of anxiety and depressive disorders amongst all students in the UK and has made it difficult for those with autism spectrum disorders (ASD) and other common mental health disorders affecting their mental wellbeing to return to face-to-face learning and on-campus life [14].

Interestingly, 53.3%, 64.4%, and 61% of all participants in year 1, 2, and 3, respectively, reported in the March 2022 (Study S3) survey that the “*lockdowns gave me a nice break from the everyday social situations*”, returning to 14.8%, 22%, and 20% for students in years 1, 2, and 3, respectively in the May 2023 (Study S4) survey (**table 19**), indicating that all students have been impacted by the social restrictions on social distancing during the Covid-19 pandemic.

We asked year 3 (level 6) students about their postgraduation career plans as bio-scientists and if these plans have changed because of the pandemic (**table 20**); data should be studied with caution as feedback was provided by different student cohorts at different time points and at different cross points during and after the pandemic. Final year students of the class 2019-2022 (yellow) showed the highest level of insecurity in reference to their postgraduation plans, while this cohort also struggled most with their pathway to resilience and adaptability to a new learning environment.

## Conclusion

This study highlights the strengths of bioscience students from all backgrounds to complete an undergraduate degree while acquiring new, graduate-ready attributes and skills, including extended IT competencies during the Covid-19 pandemic. Our results have shown that students can report poor mental health while already developing resilience, indicating tailored support can aid students’ resilience performance. Students surveyed during the Covid-19 pandemic have adjusted with ease to digital teaching provisions with current students voicing a clear preference for a subject-specific approach to teaching post-pandemic, entailing a mix of blended (synchronous) and face-to-face (on-campus) sessions, thus reducing students’ significantly risen living costs and meeting students’ often complex personal circumstances.

## Acknowledgments

Authors wish to thank all participating students 2020-2023.

## Notes

https://figshare.com/s/82d5f068b3f1c96ecc10

### Competing Interest Statement

The authors have declared no competing interest.

### Summary of Updates

To add the Supplemental Materials and to provide an external link.

## References

1. Cucinotta D, Vanelli M. WHO Declares COVID-19 a Pandemic. Acta Biomed. 2020;91(1):157–60. Epub 20200319. doi: 10.23750/abm.v91i1.9397. PubMed PMID: 32191675; PubMed Central PMCID: PMCPMC7569573.

2. UK Parliament. Coronavirus Act 2020 2020 [cited 10.01.2024]. Available from: https://www.legislation.gov.uk/ukpga/2020/7/contents/enacted.

3. Brown J, Ferguson D, Barber S. Coronavirus: The lockdown laws. Research Briefing from the House of Commons Library. 2022 [cited 10.01.2024]. Available from: https://commonslibrary.parliament.uk/research-briefings/cbp-8875/.

4. Office for Students (OfS). Letter to accountable officers: Regulation during the current phase of the coronavirus pandemic 2021 [cited 08.01.2024]. Available from: https://www.officeforstudents.org.uk/publications/letter-to-aos-regulation-during-current-phase-of-pandemic/.

5. Quality Assurance Agency for Higher Education (QAA). Subject Benchmark Statements for Bioscience 2023 [cited 04.01.2024]. Available from: https://www.qaa.ac.uk/the-quality-code/subject-benchmark-statements/subject-benchmark-statement-biosciences#.

6. Coward K, Gray JV. Audit of Practical Work undertaken by Undergraduate Bioscience Students across the UK Higher Education Sector [cited 08.02.2024]. Available from: https://www.rsb.org.uk/policy/education-policy/higher-education-policy/ug-audit-of-practical-work.

7. Gya R, Bjune AE. Taking practical learning in STEM education home: Examples from do-it-yourself experiments in plant biology. Ecol Evol. 2021;11(8):3481–7. Epub 20210203. doi: 10.1002/ece3.7207. PubMed PMID: 33898005; PubMed Central PMCID: PMCPMC8057327.

8. King CE, Trevino C, Biswas P. Online Laboratory Experiment Learning Module for Biomedical Engineering Physiological Laboratory Courses. Biomed Eng Educ. 2021;1(1):201–8. Epub 20201016. doi: 10.1007/s43683-020-00034-9. PubMed PMID: 35178535; PubMed Central PMCID: PMCPMC7567418.

9. Machuca-Tzili FA, Padilla-Ortiz AL, Martínez-Gutiérrez D. Mechanical-acoustical analogy: From laboratory to home during the COVID-19 pandemic. J Acoust Soc Am. 2023;154(3):1448–58. doi: 10.1121/10.0020828. PubMed PMID: 37675969.

10. Panebianco CJ, Iatridis JC, Weiser JR. Teaching principles of biomaterials to undergraduate students during the Covid-19 pandemic with at-home inquiry-based learning laboratory experiments. . Chem Eng Educ. 2022;56(1):22–35. Epub 20211001. doi: 10.18260/2-1-370.660-125552. PubMed PMID: 35528968; PubMed Central PMCID: PMCPMC9075043.

11. Raaper R, Brown C. The Covid-19 pandemic and the dissolution of the university campus: implications for student support practice. Journal of Professional Capital and Community. 2020;5(3-4):343–9. doi: 10.1108/JPCC-06-2020-0032.

12. Office for National Statistics (ONS). Coronavirus and the impact on students in higher education in England: September to December 2020 2020 [cited 05.01.2024]. Available from: https://www.ons.gov.uk/peoplepopulationandcommunity/educationandchildcare/articles/coronavirusandtheimpactonstudentsinhighereducationinenglandseptembertodecember2020/2020-12-21.

13. Surendran S, Hopkins S, Aji AS, Abubakar S, Clayton T, Dunuwila T, et al. Perspectives of teaching during the COVID-19 lockdown: a comparison of teaching in university bioscience programmes from around the world. Research in Science & Technological Education. 2023;41(3):1133–54. doi: 10.1080/02635143.2021.1993178.

14. Gogoi M, Webb A, Pareek M, Bayliss CD, Gies L. University Students’ Mental Health and Well-Being during the COVID-19 Pandemic: Findings from the UniCoVac Qualitative Study International Journal of Environmental Research and Public Health. 2022;19(15):9322. PubMed PMID: 10.3390/ijerph19159322.

15. Vandenbroucke JP, von Elm E, Altman DG, Gøtzsche PC, Mulrow CD, Pocock SJ, et al. Strengthening the Reporting of Observational Studies in Epidemiology (STROBE): explanation and elaboration. PLoS Med. 2007;4(10):e297. doi: 10.1371/journal.pmed.0040297. PubMed PMID: 17941715; PubMed Central PMCID: PMCPMC2020496.

16. Guetterman TC, Fetters MD, Creswell JW. Integrating Quantitative and Qualitative Results in Health Science Mixed Methods Research Through Joint Displays. The Annals of Family Medicine. 2015;13(6):554–61. doi: 10.1370/afm.1865.

17. Schoonenboom J, Johnson RB. How to Construct a Mixed Methods Research Design. Kolner Z Soz Sozpsychol. 2017;69(Suppl 2):107–31. Epub 20170705. doi: 10.1007/s11577-017-0454-1. PubMed PMID: 28989188; PubMed Central PMCID: PMCPMC5602001.

18. Cresswell JW. A concise introduction to mixed methods research. 2nd edition. International student edition. ed. Thousand Oaks, CA: SAGE Publications; 2021.

19. Tong A, Sainsbury P, Craig J. Consolidated criteria for reporting qualitative research (COREQ): a 32-item checklist for interviews and focus groups. Int J Qual Health Care. 2007;19(6):349–57. Epub 20070914. doi: 10.1093/intqhc/mzm042. PubMed PMID: 17872937.

20. Graham C. Anonymisation: Managing data protection risk - code of practice Information Commissioner’s Office: ICO; 2012 [cited 10.01.2024]. Available from: https://ico.org.uk/media/for-organisations/documents/1061/anonymisation-code.pdf.

21. UK Prime Minister’s Office. PM Commons statement on coronavirus: 22 September 2020 [cited 08.01.2024]. Available from: https://www.gov.uk/government/speeches/pm-commons-statement-on-coronavirus-22-september-2020.

22. UK Prime Minister’s Office. Prime Minister’s statement on coronavirus (COVID-19): 30 September 2020 [cited 08.01.2024]. Available from: https://www.gov.uk/government/speeches/prime-ministers-statement-on-coronavirus-covid-19-30-september-2020#:∼:text=Speech-,Prime%20Minister’s%20statement%20on%20coronavirus%20(COVID%2D19)%3A%2030%20September,at%20the%20coronavirus%20press%20conference.&text=and%20I%20explained%20that%20the,expect%20many%20more%20daily%20deaths.

23. UK Government - Department for Education. Christmas guidance set out for university studentslJ 2020 [cited 08.01.2024]. Available from: https://www.gov.uk/government/news/christmasguidance-set-out-for-university-students.

24. Hubble S, Bolton P, Lewis J. Coronavirus: HE/FE return to campus in England 2021. Research Briefing from the House of Commons Library.: UK Parliament; 2021 [cited 10.01.2024]. Available from: https://commonslibrary.parliament.uk/research-briefings/cbp-9142/.

25. Office for National Statistics (ONS). Economic activity status by ethnic group 2023 [cited 08.02.2024]. Available from: https://www.ons.gov.uk/datasets/RM018/editions/2021/versions/3.

26. National Office for Statistics (ONS). International migration, England and Wales. 2022 [updated 02.11.2022cited 25.01.2024]. Available from: International migration, England and Wales: Census 2021.

27. UK Ministry of Housing, Communities & Local Government,. English indices of deprivation 2019 [cited 24.01.2024]. Available from: https://www.gov.uk/government/statistics/english-indices-of-deprivation-2019.

28. Office for National Statistics (ONS). Coronavirus and higher education students: England, 24 May to 2 June 2021 2021 [cited 17.01.2024]. Available from: https://www.ons.gov.uk/peoplepopulationandcommunity/healthandsocialcare/healthandwellbeing/bulletins/coronavirusandhighereducationstudents/england24mayto2june2021.

29. Office for Students (OfS). The National Student Survey: Student experience during the pandemic 2021 [cited 17.01.2024]. Available from: https://www.officeforstudents.org.uk/publications/the-national-student-survey-student-experience-during-the-pandemic/.

30. Bashir A, Bashir S, Rana K, Lambert P, Vernallis A. Post-COVID-19 Adaptations; the Shifts Towards Online Learning, Hybrid Course Delivery and the Implications for Biosciences Courses in the Higher Education Setting. Frontiers in Education. 2021;6. doi: 10.3389/feduc.2021.711619.

31. Ortega-Donaire L, Bailén-Expósito J, Álvarez-García C, López-Medina IM, Álvarez-Nieto C, Sanz-Martos S. Satisfaction of Online University Education during the COVID-19 Pandemic. Healthcare (Basel). 2023;11(10). Epub 20230514. doi: 10.3390/healthcare11101421. PubMed PMID: 37239706; PubMed Central PMCID: PMCPMC10218284.

32. Albiser E, Echazarra A, Fraser P, Fülöp G, Schwabe M, Tremblay K. School education during Covid-19: Were teachers and students ready? : OECD; 2020 [cited 17.01.2024]. Available from: https://www.oecd.org/education/United-Kingdom-coronavirus-education-country-note.pdf.

33. McManus S, Bebbington P, Jenkins R, Brugha T, (eds). Mental Health and Wellbeing in England: Adult Psychiatric Morbidity Survey 2014 Leeds: NHS digital; 2016 [cited 18.01.2024]. Available from: https://assets.publishing.service.gov.uk/media/5a802e2fe5274a2e8ab4ea71/apms-2014-full-rpt.pdf.

34. Anandaciva S, and the The King’s Fund. How does the NHS compare to the health care systems of other countries? 2023 [cited 10.01.2024]. Available from: https://www.kingsfund.org.uk/publications/nhs-compare-health-care-systems-other-countries?page=1#comments-top.

35. OECD. Health at a Glance 2021 [cited 12.01.2024]. Available from: https://www.oecd-ilibrary.org/content/publication/ae3016b9-en.

36. Chen T, Lucock M. The mental health of university students during the COVID-19 pandemic: An online survey in the UK. PLoS One. 2022;17(1):e0262562. Epub 20220112. doi: 10.1371/journal.pone.0262562. PubMed PMID: 35020758; PubMed Central PMCID: PMCPMC8754313.

37. Office for National Statistics (ONS). Coronavirus and higher education students: 25 February to 7 March 2022 [cited 30.01.2024]. Available from: https://www.ons.gov.uk/peoplepopulationandcommunity/healthandsocialcare/healthandwellbeing/bulletins/coronavirusandhighereducationstudents/25februaryto7march2022.

38. Montacute R, and the Sutton Trust. Social mobility and Covid-19 - Implications of the Covid-19 crisis for educational inequality 2020 [cited 29.01.2024]. Available from: https://www.suttontrust.com/wp-content/uploads/2020/04/COVID-19-and-Social-Mobility-1.pdf.

39. Office for National Statistics (ONS). Deprivation inequalities in the experiences of GCSE students during coronavirus (COVID-19), England: September 2021 to March 2022 2023. Available from: https://www.ons.gov.uk/peoplepopulationandcommunity/educationandchildcare/articles/deprivationinequalitiesintheexperiencesofgcsestudentsduringcoronaviruscovid19england/september2021tomarch2022.

40. Di Pietro G. The impact of Covid-19 on student achievement: Evidence from a recent meta-analysis. Educational Research Review. 2023;39. doi: 10.1016/j.edurev.2023.100530.

41. National Office for Statistics (ONS). Cost of living and higher education students, England: 30 January to 13 February 2023 [cited 30.01.2024]. Available from: https://www.ons.gov.uk/peoplepopulationandcommunity/educationandchildcare/bulletins/costoflivingandhighereducationstudentsengland/30januaryto13february2023.

42. Salem IE, Al-Alawi A, Moosa S, El-Maghraby L, Alkathiri NA, Elbaz AM. Examining different learning modes: A longitudinal study of business administration students’ performance. International Journal of Management Education. 2024;22(1). doi: 10.1016/j.ijme.2023.100927.

