## Supplements S1-S4 for "*Learn!Bio* - A time-limited cross-sectional study on biosciences students’ pathway to resilience during and post the Covid-19 pandemic at an UK university from 2020-2023 and insights into future teaching approaches"

**Short title:**

*Learn!Bio – Bioscience students pathway to resilience during and post the Covid-19 pandemic*

Katy Andrews^1#^, Rosalie Stoneley^1#^, and Katja Eckl^1*^

Department of Biology, Edge Hill University, L39 4QP Ormskirk, Lancashire, United Kingdom.

^#^ authors contributed equally.

**Supplements S1-S4**

**Supplements S1**

Survey questionnaires for studies 1-4 can be downloaded from Edge Hill University’s data depository at <https://figshare.com/s/82d5f068b3f1c96ecc10>

**Supplement S2**

Table S2: Students’ accommodations in semester 1, academic year 20/21

| Where to you live this semester? | **Study 1 (Nov 20)** | | |
| --- | --- | --- | --- |
|  | **L4** | **L5** | **L6** |
| On campus | **34.6** | 10.7 | **25.0** |
| Private off-campus | 19.2 | **25.0** | **31.3** |
| at home (parents) | 26.9 | 42.9 | **25.0** |
| at home (own/with partner/with children) | 19.2 | 17.9 | 18.8 |
| prefer not to say | 0.0 | 0.0 | 0.0 |

Explanations: Responses from the *Learn!Bio* Study 2 in April 2021, one month after the third national lockdown had come to an end. Multiple answers possible. Numbers in brackets after the level L4-L6 indicate the total participants followed by total answers provided. Results displayed in percentages.

**Supplement S3**

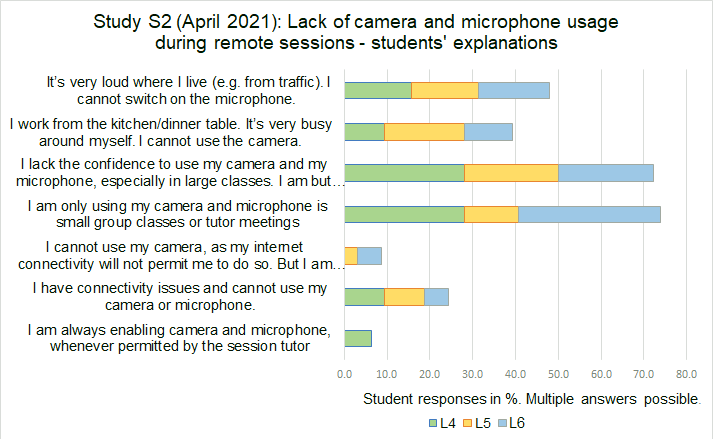

**Figure S3:** Lack of microphone and camera use during online classes. Students’ responses to a multiple-answers question in the April 2021 study S2. For year 1 (level 4, L4), 16 participants provided 32 answers, for level 5 (L5) 11 participants provided 32 answers, and for level 6 (L6), 7 participants provided 18 answers. Results displayed in percentages.

**Supplement S4**

Table S4: New skills and talents developed during the Covid-19 pandemic.

| ***Looking back at the past academic year: Do you believe you have developed or nurtured a wealth of skills or talents as a consequence of long times spent in lockdown? Please indicate if any of the below apply to you.*** | **Study 2 (Apr 21)** | | |
| --- | --- | --- | --- |
|  | **L4 (16/96)** | **L5 (11/34)** | **L6 (7/37)** |
| I am now (much more) computer and IT literate | 50.0 | 27.3 | 71.4 |
| I can fix many of the standard IT/internet/Microsoft Office problems myself | 62.5 | 36.4 | 71.4 |
| I am able to use frequently used conference software tools, including Blackboard Collaborate, MS Teams, and Zoom | 87.5 | 72.7 | 57.1 |
| I am more relaxed about smaller problems and hurdles - It is the whole picture which counts | **50.0** | **9.1** | **28.6** |
| I have learnt to skim read scientific books and papers– quickly skimming lots of literature in shortest time and still understanding its content. | 68.8 | 36.4 | 85.7 |
| I am much better with MS Office, including PowerPoint, Excel and Word. | 75.0 | 54.5 | 57.1 |
| I know how to seek remote support (e.g. from the department admin team, my personal tutor, or the Catalyst team) | 68.8 | 45.5 | 85.7 |
| I have learned to work self-paced towards a deadline and to plan ahead in a timely fashion | **56.3** | **27.3** | **71.4** |
| I can work remotely/virtually in teams, conducting group activities and group assessments. | **75.0** | **0.0** | **0.0** |

Explanations: Responses from the *Learn!Bio* Study 2 in April 2021, one month after the third national lockdown had come to an end. Multiple answers possible. Numbers in brackets indicate the total participants followed by total answers provided. Results displayed in percentages.
